# Correlated evolutionary rates reveal novel components and cross-compartment connectivity in plant proteostasis systems

**DOI:** 10.1101/2024.08.22.609246

**Authors:** Tony C. Gatts, Elizabeth A. Rehmann, Daniel B. Sloan, Evan S. Forsythe

## Abstract

Plant cells rely on an interconnected network of proteins interacting at many levels (*e*.*g*. physical enzyme complexes, gene regulatory modules, and biosynthetic pathways). Pairs of proteins that interact at any of these levels have been shown to exhibit phylogenetic signatures of evolutionary rate covariation (ERC), providing a basis for detecting functional interactions among proteins. Here, we apply *ERCne*t, a bioinformatic tool for performing genome-scale ERC analyses, to predict a plant protein-protein interactome network. We find a clustered set of proteins that exhibit strong signatures of ERC with the plastid caseinolytic protease (Clp) and other plastid proteostasis components, thus forming a functional module within the network. In addition to including proteins with known or predicted functions in protein import, transcription, translation, and degradation in plastids, the module also includes proteins with previously unknown molecular function, thus providing evidence that these proteins may contribute to plastid proteostasis in novel ways. Perhaps the most surprising members of this module are a set of proteins that are not thought to localize to the plastid at all. These proteins include a mitochondrial-localized pentatricopeptide repeat (PPR) protein with genetic evidence of interaction with the mitochondrial Clp system and two nuclear-localized actin-related proteins involved in chromatin remodeling and epigenetic regulation of nuclear genes. We speculate that these non-plastid-localized proteins act as mediators of organellar crosstalk and retrograde signaling of cellular proteostasis status in plants. In summary, our results highlight the connected nature of plant proteostasis systems and point to a promising set of novel proteostasis protein candidates.

## Introduction

Every cellular process relies on finely tuned interactions among gene products (Hidalgo et al. 2007; Yu et al. 2008b; Wright et al. 2024). This division of labor is especially pronounced in cytoplasmic organelles with their own genomes (mitochondria and plastids) whose functions depend on interactions between organellar gene products and proteins that are encoded in the nucleus, translated in the cytosol, and imported into the organelle. These proteins undergo folding and are incorporated into complexes, often in direct physical contact with organellar proteins (i.e. cytonuclear complexes) (Rand et al. 2004; Greiner and Bock 2013; Barreto et al. 2018; Sloan et al. 2018).

Communication between cellular compartments is required for plant growth (Woodson and Chory 2008) and response to stress (Llamas and Pulido 2022; Lemke and Woodson 2023). However, the mechanisms underlying retrograde (organelle-to-nucleus) and anterograde (nucleus-to-organelle) signaling are not fully resolved (Richter et al. 2023). Complex regulation is also required to maintain proper protein accumulation levels in plastids (plastid proteostasis). Many overlapping factors contribute to plastid proteostasis, including (1) the rate of plastid transcription and translation, (2) the rate of transcription, translation, targeting, import, and folding of nuclear-encoded proteins that function in plastids (N-pt proteins), and (3) the quality control and degradation of both plastid-encoded and N-pt proteins (Paila et al. 2015; Llamas and Pulido 2022; Van Wijk 2024). Moreover, all of these processes must be able to respond to external stimuli, including photodamage, which is especially pronounced in plastids given their role in light harvesting (Kato and Sakamoto 2018). Mutations affecting genes in these pathways can yield characteristic plastid proteostasis mutant phenotypes (Park and Rodermel 2004; Yu et al. 2008a; Wang et al. 2018), highlighting the degree of interconnectivity between the disparate components of plastid proteostasis.

Many of the most well-studied functional connections in plastid proteostasis systems include direct physical protein-protein interactions within heteromeric enzyme complexes (Nishimura and van Wijk 2015). However, pairs of proteins that do not physically interface or even localize to the same cellular compartment can also be functionally related based on their contribution to overlapping processes, constituting a more cryptic level of interaction (Clark et al. 2012; Forsythe et al. 2021; Little et al. 2024). These “co-functional” interactions are not detectable with conventional biochemistry-based methods, such as coimmunoprecipitation (Hidalgo et al. 2007). One hallmark of co-functional interactions (regardless of whether they involve physical contact) is the tendency for pairs of proteins to exhibit correlated rates of sequence evolution across species, which can be detected with phylogenetic analyses that measure evolutionary rate covariation (ERC). Analyses of ERC have been performed in bacteria (Ramani and Marcotte 2003), and yeast (Clark et al. 2012; Steenwyk et al. 2022), mammals (Goh et al. 2000; Priedigkeit et al. 2015), insects (Findlay et al. 2014; Yan et al. 2019; Tao et al. 2024). More recently, these analyses have been applied to plants (Forsythe et al. 2021; Gatts et al. 2024), despite challenges due to their rampant gene and genome duplication (Wendel 2015; Panchy et al. 2016).

Because ERC detects shifts in evolutionary rate, the method holds the most statistical power in cases in which species vary substantially in gene-specific rates of evolution. Despite the importance of the plastid proteostasis machinery and general plastid function in photosynthetic plants, the plastid genomes (plastomes) of several angiosperm species have undergone dramatically accelerated evolution (Guisinger et al. 2008; Sloan et al. 2014; Williams et al. 2019). These acceleration events occurred independently several times across the angiosperm phylogeny but the evolutionary mechanisms that cause these punctuated periods of rapid evolution, despite a background slow rate of the evolution in most species, remain a mystery. The repeated, parallel nature of these shifts presents an opportunity to apply ERC to detect co-functional genes that underwent correlated rate shifts. We previously applied ERC in angiosperms to show that a subset of N-pt genes exhibit correlated evolution with plastomes during these rate accelerations (Forsythe et al. 2021). We found that this subset of N-pts was enriched for genes associated with plastid proteostasis, suggesting that perturbation of proteostasis systems may be involved in the initial events that precipitated the mysterious rate accelerations. The plastid caseinolytic protease (Clp) complex shows a particularly dramatic level of rate acceleration, which is consistent with the idea that rate acceleration is tied to plastid proteostasis. It is possible that Clp and other proteases play a role in integrating signals from the disparate components of the proteostasis apparatus (Llamas and Pulido 2022; Van Wijk 2024). However, previous plastome-vs-nucleus (one-vs-all) analyses lacked the dimensionality and resolution to fully reveal the functional interactions in plastid proteostasis systems. Extending ERC analyses to infer interactome networks could reveal clustered groups of proteins (i.e. network communities) with related functions, indicative of functional modules in the organization of plant cellular systems (Ahn et al. 2010; Tripathi et al. 2019).

Here, we predict an ERC-based protein-protein interactome at a genome-wide all-by-all scale for the first time in plants. We reveal an interaction network comprised of 2,951 proteins connected via 3,228 predicted pairwise interactions. This network shows significant clustering of functionally related genes at both local and global scales. We identify a region of the network that shows strong evidence of being a functional module that is highly enriched for plastid-encoded and N-pt proteins but also contains a small number of nuclear proteins not previously known to function in plastids. We use our network structure to gain insight into plastid proteostasis evolution by focusing on known members of the Clp complex, and we identify nine candidates as novel Clp interactions partners, including proteins likely involved in pathways that contribute to plastid proteostasis (e.g., plastid import, transcription, and translation). These candidates that exhibited strong ERC with Clp also include proteins with unknown function and, surprisingly, proteins that localize to the nucleus, cytoplasm, and mitochondrion. We discuss how our genome-wide interactome network points to evidence of cross-compartment co-evolution of proteostasis systems.

## Results

### Predicting the plant protein interactome from evolutionary signatures

We made use of proteome sequences from select angiosperm species that were previously compiled to study plastid-nuclear co-evolution (Forsythe et al. 2021). We analyzed this dataset using the recently developed *ERCnet* program for large-scale phylogenomic prediction and network analyses in protein-protein interactomes (Gatts et al. 2024). *ERCnet* filters yielded 8,118 gene families suitable for ERC analysis. These gene families included nuclear-encoded proteins in addition to seven gene families for ‘plastome-partitions’, which represent previously analyzed groupings of functionally related plastid-encoded proteins (Forsythe et al. 2021). We performed 30.8 million of the possible 32.9 million pairwise ERC comparisons between the 8,118 gene families, after excluding possible comparisons that did not have sufficient overlap in taxon sampling (Fig. 1). These results were filtered for significant positive correlations (hereafter referred to as ‘ERC hits’), yielding an interactome network (Fig. 2) composed of 3,228 ERC hits (edges in the network) involving a total of 2,951 proteins (nodes in the network). We found that known members of the Clp complex are tightly clustered within the network and are connected by thick edges (Fig. 2), suggesting widespread and strong ERC signatures within the Clp complex. This result is consistent with previous analyses that showed ERC between ClpP1 and N-pt Clp subunits (Williams et al. 2019; Forsythe et al. 2021) as well as among N-pt Clp subunits (Rei Liao et al. 2022).

**Fig. 1:**
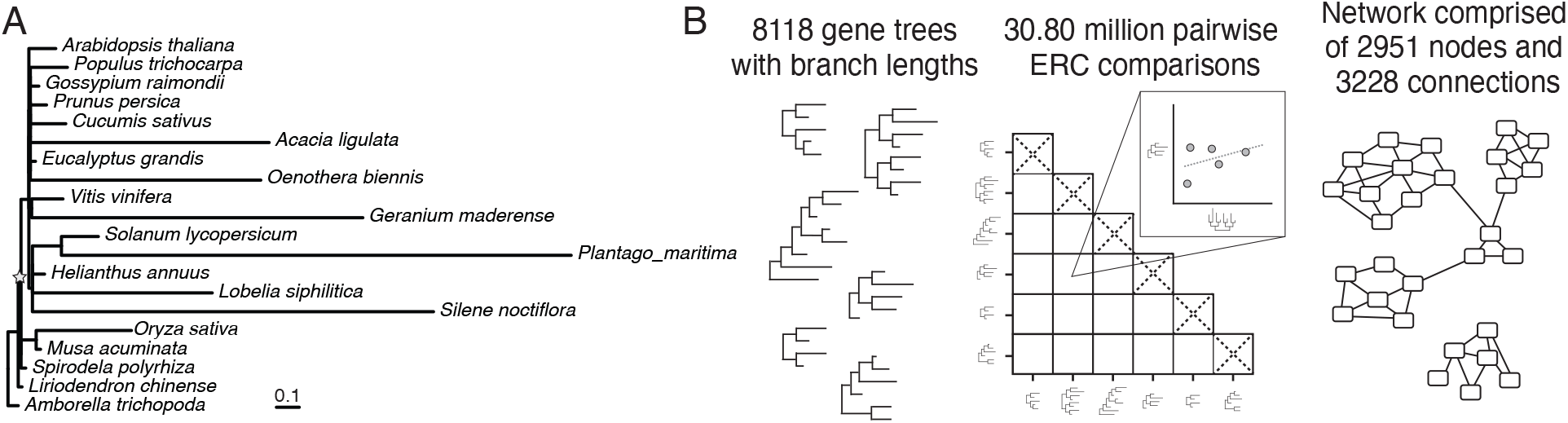
ERC analyses of all pairwise combinations of protein-protein interactions in angiosperms. (A) Phylogenetic tree displaying twenty angiosperm species included in this analysis. Branch lengths were inferred from the ClpP1 protein sequence. Star indicated the node that was used as the ‘ingroup’ node, to the exclusion of the two outgroup species used in this study, *Amborella trichopoda* and *Liriodendron chinense*. Figure adapted from (Forsythe et al. 2021). (B) Summary of the major analytical steps of *ERCnet* that were applied to the angiosperm dataset.

**Fig. 2:**
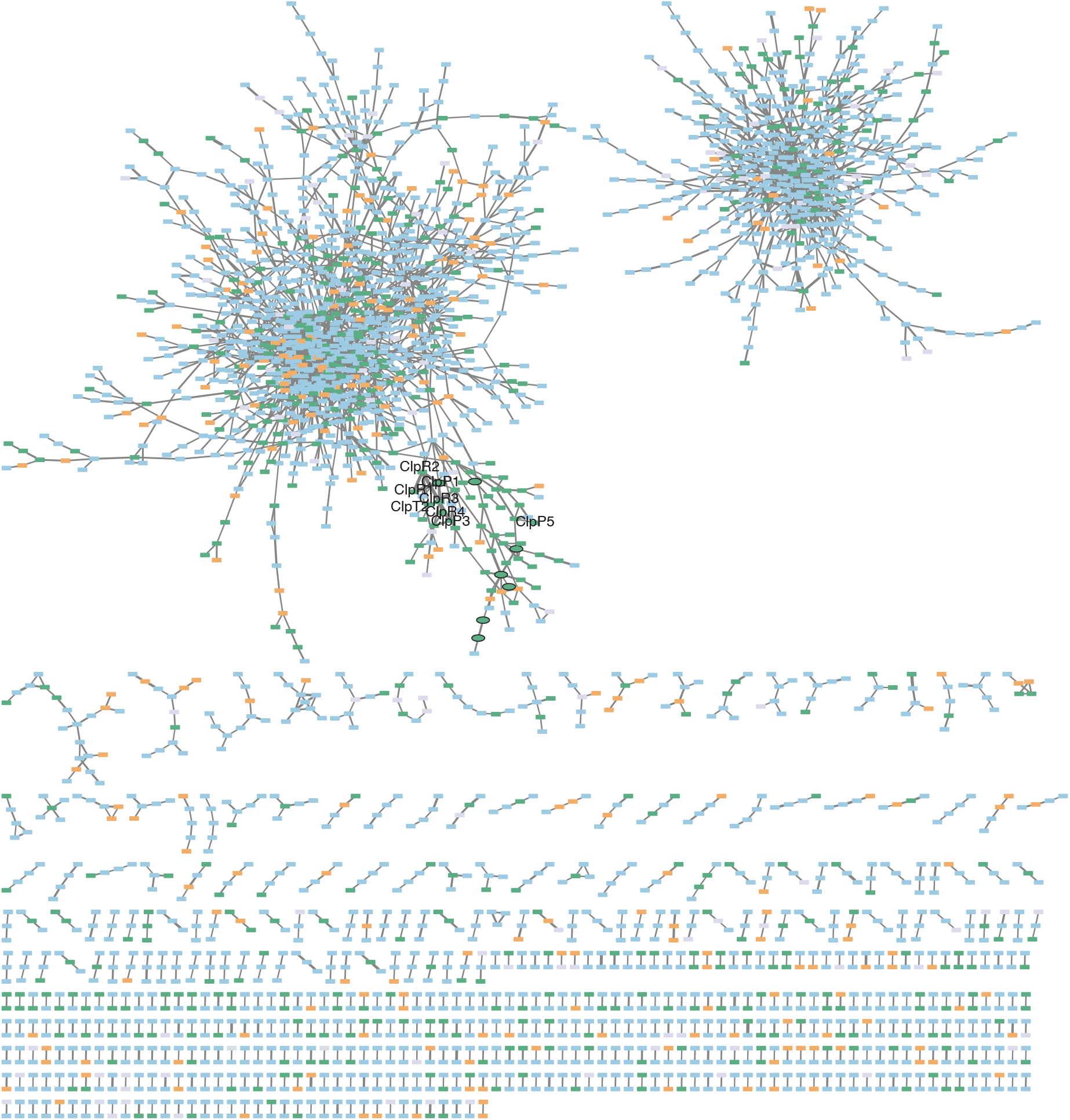
A global ERC network. Interaction network displaying strong ERC hits (Pearson p<0.0001, Spearman p<0.0001, and R^2^>0.5). Plastid-encoded partitions are displayed as ovals. Nuclear-encoded proteins are displayed as rectangles. Points are colored by predicted subcellular localization (CyMIRA; Forsythe et al., 2019). Green: plastid-localized; orange: mito-localized; blue: other localization; gray: localization unknown. Connection edge weight is scaled by Pearson p-value. Known Clp subunits are labeled. Plot prepared with *Cytoscape*.

### Functional clustering within the interaction network

To ask whether the network contains functional clustering information that goes beyond the Clp complex, we inspected its structure to identify groups of interconnected nodes, referred to as ‘communities’ in graph theory (Radicchi et al. 2004). We used *Cytoscape* to identify communities of interest within in the network and applied gene ontology (GO) analyses to ask whether certain types of proteins are enriched within these communities (see Methods). First, we identified the network community that contains all the Clp complex proteins (Fig. 2). This community contains a total of 83 nodes. Among these nodes are all seven plastome partitions, suggesting the community may be clustered on the basis of broader plastid function. Consistent with this idea, we found that the remaining nuclear-encoded nodes are highly enriched for being plastid-localized (Fig. 3A). These genes also show modest but significant enrichment for being involved in translation. These enrichment results provide evidence that this community constitutes a module of co-functional proteins involved in plastid functions, the members of which could yield important insights into novel plastid-nuclear interactions (see below). We also performed enrichment analyses on the largest community in the network, which includes 1033 nodes and has a small number of connections to the plastid community described above. For this community we did not find any significantly enriched GO categories (Fig. 3B). However, it is possible that its smaller sub-communities and pairwise connections may still contain important functional information. Finally, we tested a third community with 483 nodes that is completely disconnected from the rest of the network and found that it showed enrichment for GO categories associated with transcription factor activity (Fig. 3C), which could suggest this community is of interest as a co-regulatory module. Given the strength of the enrichment results for the 83-node community, we focus our detailed analysis of individual genes on the members of this community (see Table 1, Fig. 5, and Discussion).

**Table 1:**
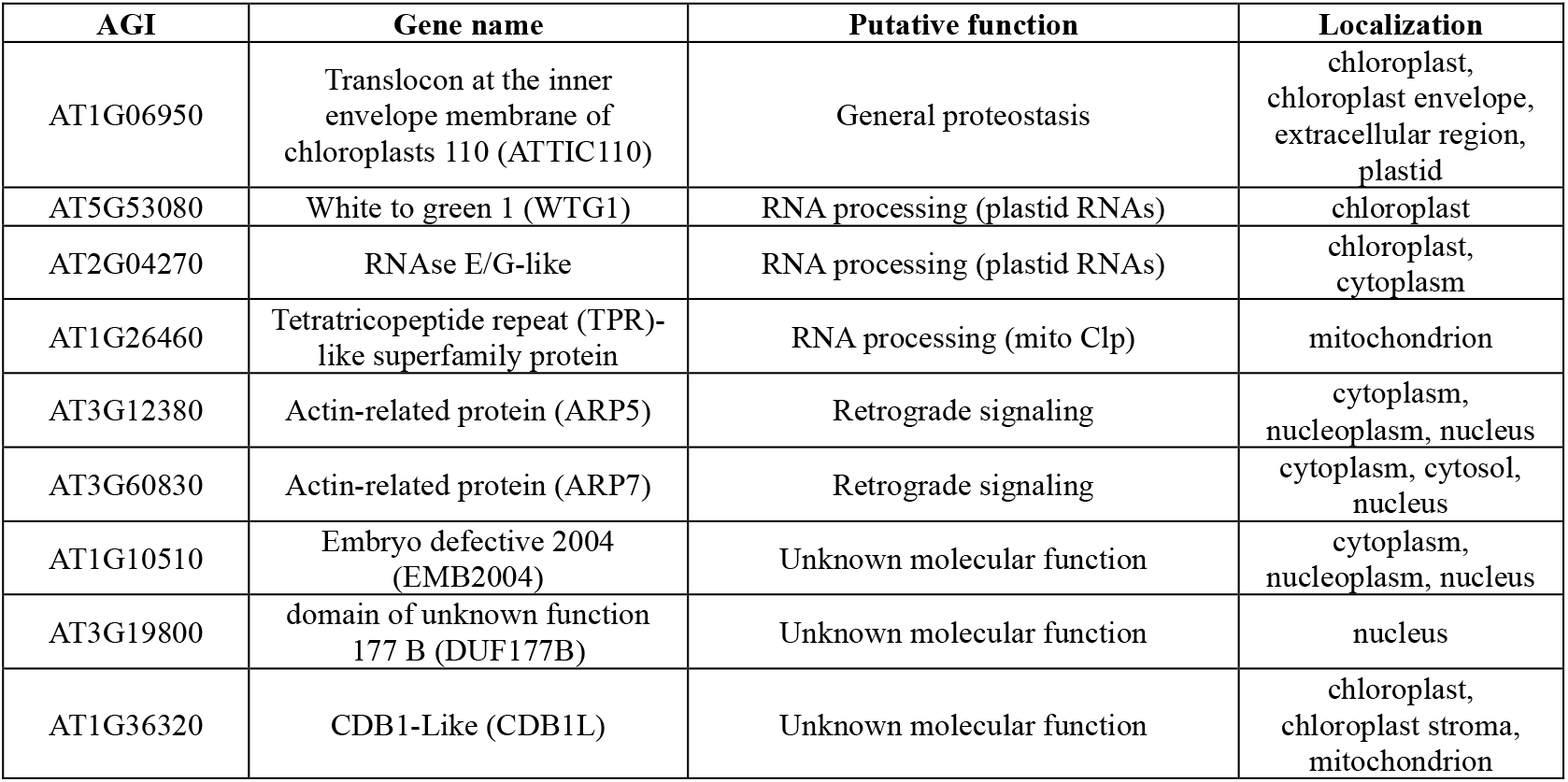
Novel candidate interactors with the Clp proteostasis system. Candidate interactors with the Clp proteostasis system. Proteins with two or fewer degrees of separation from ClpP1 in the interaction network are listed. Previously known Clp subunits are excluded for brevity.

**Fig. 3:**
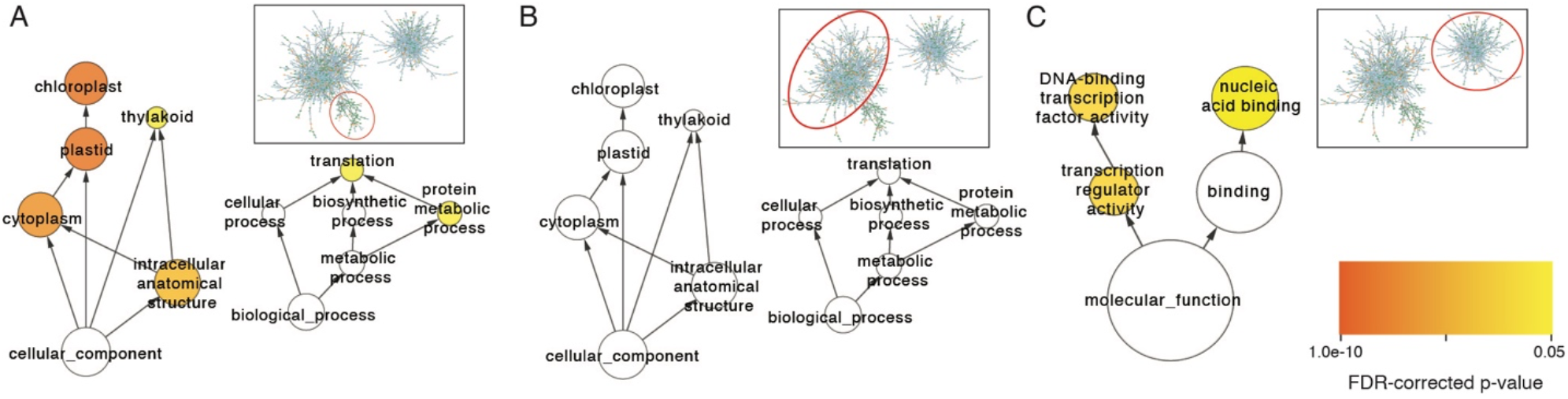
Functional enrichment in network communities. GO categories enriched in network communities. Enriched categories are plotted as a network displaying hierarchical GO category structure using the *BINGO* plugin for *Cytoscape*.

In addition to testing for functional enrichment at the level of specific communities or complexes, we also asked whether functionally related proteins tend to cluster at the level of the full network. To test for global pattens of functional clustering, we calculated the assortativity coefficient, a network summary statistic that quantifies assortative (like clustered with like) and disassortative (like clustered with unlike) clustering of nodes in a network based on node attributes (Newman 2003). We first assigned attributes to the nodes in the network based on subcellular localization information obtained from *Arabidopsis thaliana* (Forsythe et al. 2019) and found assortative (positive) clustering (assortativity coefficient=0.068; p=6.81e-8) (Fig. 4A). This result indicates that the network successfully clusters colocalized proteins and that this signature is detectable across the full network.

**Fig. 4:**
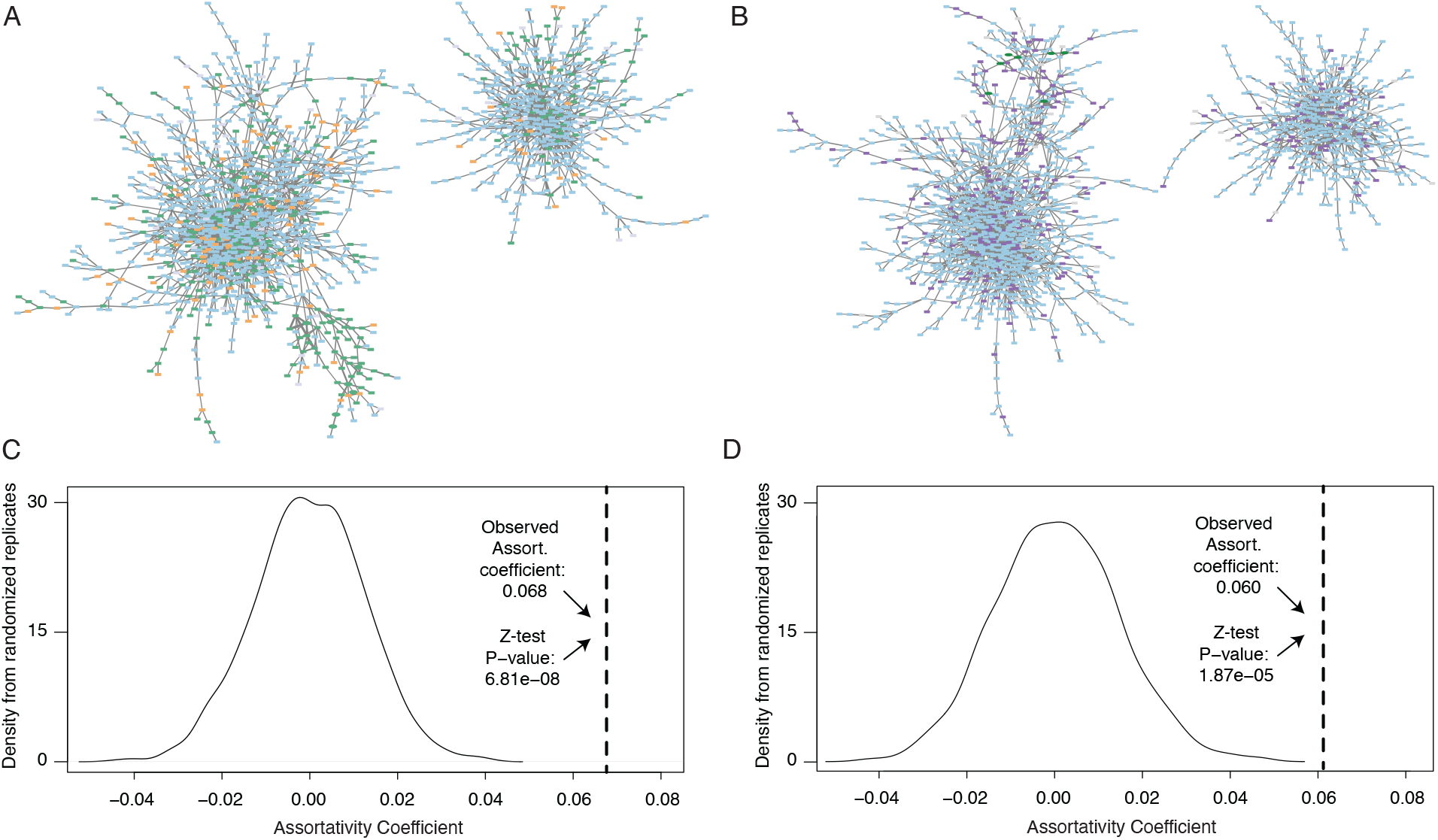
Functional clustering in the network. Network-wide assortative clustering measured with assortativity coefficient. (**A and C**) Network color scheme and assortativity calculations based on subcellular localization. (**B and D**) Network color scheme and assortativity calculations based on a network of proteostasis genes predicted through a combination of genome-wide co-expression analyses and a manually curated set of peptidase and peptidase-interacting proteins (Majsec et al., 2017; Fig. 4). Purple nodes are predicted members of the Majsec et al., network and blue nodes are all other genes. Assortativity coefficient was calculated using the *R igraph* package. Null distributions were obtained from 1000 replicates in which node attributes were randomized across the network.

Next, we used assortativity to ask if the interactions predicted in our ERC-based network are consistent with networks predicted with an orthogonal method, co-expression. A previous study used created a co-expression network of *A. thaliana* proteins and used this network to identify the genome-wide set of plastid and mitochondrial proteolysis genes (97) and co-expressed genes (1645) (Majsec et al. 2017). We identified these genes within our ERC-based network and used them as node attributes to perform an assortativity test asking if our network successfully clusters them. We found that assortativity analysis of these node attributes also yielded significantly positive clustering (assortativity coefficient=0.060; p=1.87e-05). This result suggests that the ERC-based network shows significant overlap with a co-expression-based network for the predicted proteostasis proteins. Taken together, the community enrichment and functional clustering results indicate that the connections in our network are biologically informative. These results provide an encouraging validation of the utility in using ERC-based network to identify co-functional proteins involved in proteostasis and justify detailed analyses of the predicted protein interactions.

### Network structure within the plastid functional module

Given the above signatures of functional interactions exhibited in the network and the observation that these signatures appear to be especially strong in relation to plastid-nuclear interactions, we focused on the 83-node plastid-enriched community (Fig. 5A). The structure of this community shows that a set of nodes representing plastid-encoded proteins (concatenated subunits of photosynthesis enzyme complexes, concatenated subunits of the plastid-encoded RNA polymerase (PEP), concatenated subunits of the plastid ribosome, the AccD subunit of the heteromeric acetyl-CoA carboxylase complex, and the intron maturase MatK) are directly connected to each other, with the exception of the linkage between the ribosomal protein node and the MatK node, which is an indirect connection via a mitochondrial-localized nuclear-encoded protein (ribosomal protein MS85) (Fig 5B). The remaining two plastid partition nodes (ClpP1 and Ycf1/2) are located in a more distant region of the community ≥ 4 connections from the other plastid partitions. ClpP1 and Ycf1/2 are separated from each other by three edges, meaning they exhibit less direct connection than the other plastid-encoded nodes.

**Fig. 5:**
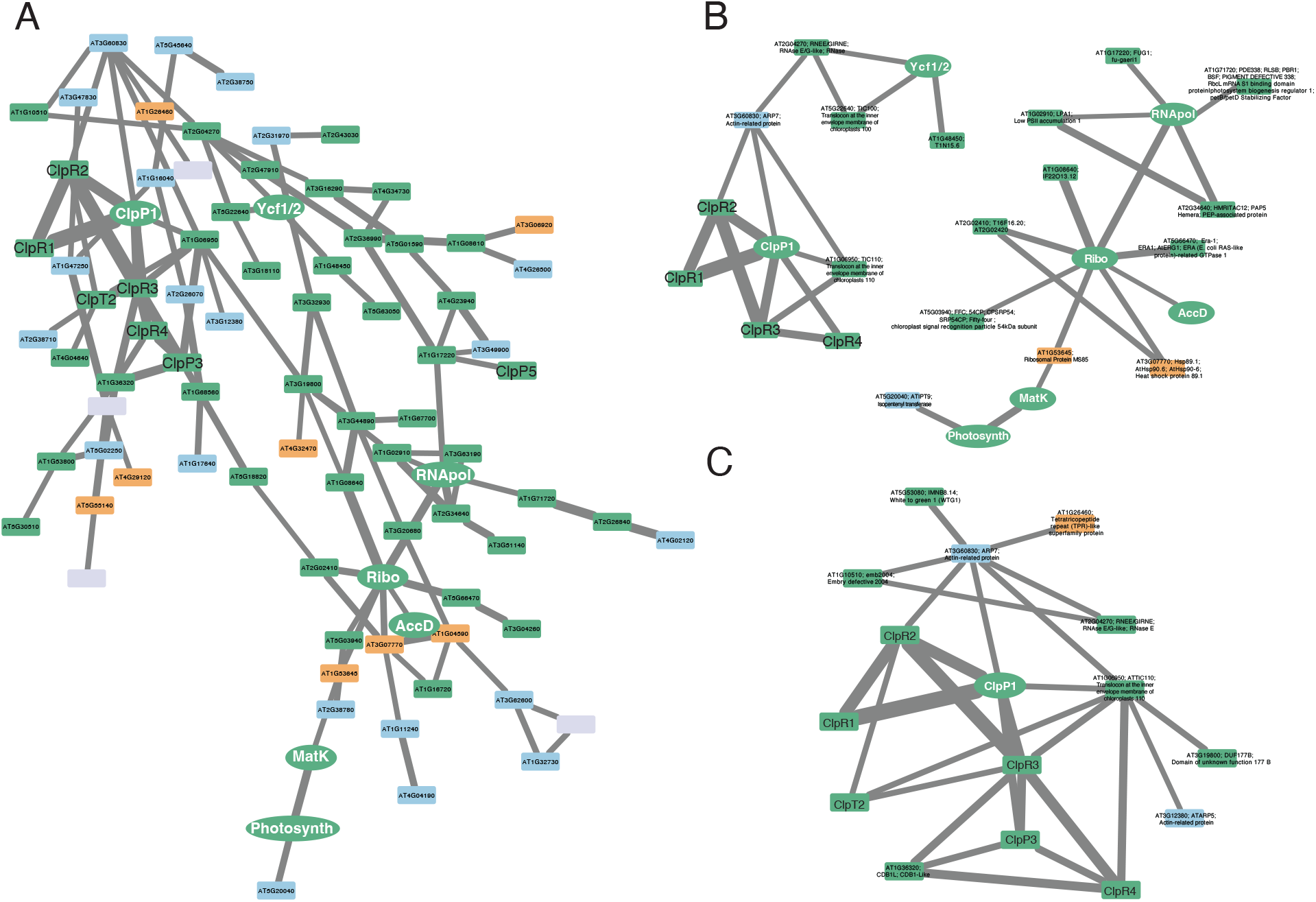
Co-functional interactions in the plastid interactome. Subgraphs containing plastid and plastid-localized proteins. Node shape and color are the same as Fig. 2. (**A**) The full community that contains plastid partitions. *Arabidopsis* Genome Initiative (AGI) identifiers are shown for nodes in which *A. thaliana* was present in the gene family. (**B**) Proteins involved in direct interactions with a plastid partition. (**C**) Proteins involved 1^st^-degree (direct) or 2^nd^-degree interactions with ClpP1. (**B and C**) Gene descriptions from TAIR are shown for each protein.

Given the signs of ERC driven by plastid proteostasis, we zoomed in on the nodes surrounding the Clp complex; we define the nodes separated by ≤ 2 edges from ClpP1 as our ‘short-list’ of Clp iteration partners (Fig. 5C and Table 1). Notably, all analyzed Clp proteins are included in this short-list, except for ClpP5, which, instead, is indirectly connected to the RNA polymerase plastome partition via the N-pt protein FUG1 (AT1G17220) translation initiation factor. Beyond the six known Clp proteins, ClpP1 is connected to nine additional proteins which are not designated to be part of the Clp complex (Table 1). These include proteins putatively involved in plastid protein import, plastid RNA processing, mitochondrial RNA processing as well as actin-related proteins and proteins with unknown function. Each member of this shortlist represents a putative novel interaction with the Clp complex and plastid proteostasis systems. The results presented here exhibit how ERC can be leveraged to yield networks with functional information spanning local to global levels of organization of the plant protein-protein interactome.

## Discussion

### Correlated evolution among physical and co-functional interactors

There is growing evidence that a classical view of compensatory evolution at physical interaction interfaces (De Juan et al. 2013) is not the sole force that drives patterns of correlated evolution (Little et al. 2024). Prior work has demonstrated ERC between enzymes that catalyze separate steps of a biosynthetic pathway without physically contacting each other (Clark and Aquadro 2010; Clark et al. 2012) and even between proteins that localize to different cellular compartments (Forsythe et al. 2021). Little *et al*. (2024) explicitly tested the relative contributions of physical vs non-physical forces and found that ERC hits tend to be a mixed bag composed of both types of interaction partners. Our results add to the notion that ERC signatures arise due to many types of interactions. Building on previous results that identified ERC between plastid-localized and cytosolic proteins (Forsythe et al. 2021), we find strong ERC signatures that indicate co-functional interaction of plastid proteins with mitochondrial-localized and nuclear-localized proteins, representing an even larger degree of physical separation that spans at least two membrane systems (Table 1). These results expand our definition of ‘functional interactions’, which opens the door to discovering critical non-physical interactions that may have been previously overlooked.

### Proteostasis is at the intersection of multiple plastid pathways

The plastid proteostasis system at-large is emerging as one of the most striking examples of ERC created by co-functional interactions, including interactions that span cellular compartments (Forsythe et al. 2021). Monitoring and communicating proteostasis status across different cellular compartments and coordinating the action of multiple interleaving pathways may be critical. One logical candidate for a junction point in a proteostasis network would be plastid protein import machinery. Plastid import not only impacts protein levels inside of plastids, but also impacts the cytosolic accumulation of potentially toxic unfolded pre-proteins *en route* to plastid import (Paila et al. 2015). Consistent with this idea, our network indicates that Translocon at the Inner Envelope Membrane of Chloroplasts 110 (TIC110; AT1G06950) shows a high degree of ERC signature with the Clp system. TIC110 is a well-characterized component of the protein import complex at the inner membrane of plastids (Inaba et al. 2005). TIC110 is present from glaucophytes and red algae to flowering plants and is almost always encoded by a single gene (Tsai et al. 2013). It has been postulated that Clp could act as a quality control system that directly screens proteins as they are imported into plastids and that Clp may even contribute to the protein import motor (Paila et al. 2015). Consistent with this hypothesis, HSP93/ClpC has been described as both a member of the import apparatus and an accessory protein in the Clp complex (Kovacheva et al. 2005; Sjögren et al. 2014; Flores-Pérez et al. 2016; Huang et al. 2016; Yuan and Van Wijk 2024). Our results offer additional evidence connecting the Clp complex to protein import and indicate that TIC110 may play a central role in mediating interaction at either a functional or physical level between Clp and TIC, which is consistent with observations that TIC110 acts as a scaffold for protein-protein interactions (Paila et al. 2015). Moreover, the location of TIC110 in our network, connected to seven other nodes, may indicate the TIC110 is an important player at the junction of many intersecting proteostasis pathways. It should be noted that the HSP93/ClpC gene family did not survive the *ERCnet* filters, meaning we could not probe whether it also exhibits ERC with TIC110.

Interestingly, the connected ClpP1 and TIC110 nodes in the network are located just three edges from another pair of connected nodes corresponding to the plastid Ycf1/2 and TIC100 (AT5G22640), another member of the import complex (Fig. 5B). This interaction is notable because Ycf1 (alternatively named TIC214) was found to be part of the TIC complex and Ycf2 is related to members of the FtsH protease complex (Kikuchi et al. 2018), representing yet another example of ERC between genes involved in import and proteostasis. This observation could be a sign that the evolutionary forces that connect Clp and TIC110 have acted in parallel for Ycf1/Ycf2 and TIC100.

### RNA processing and a possible connection between plastid and mitochondrial Clp systems

There is evidence that at least three of the nine proteins in our short-list of ERC hits (AT5G53080, AT2G04270, and AT1G26460) are involved in organellar RNA processing steps that are critical to organelle biogenesis, suggesting a tie between RNA processing and plastid proteostasis. Post-transcriptional regulation is extensive in plastid genes; for example, changing light conditions lead to altered rates of plastid RNA degradation (Danon and Mayfield 1994) and translation (Salvador and Klein 1999). Organellar gene expression appears to rely more heavily on post-transcriptional regulation than does nuclear gene expression (Small et al. 2013; Chotewutmontri and Barkan 2018), which could possibly explain why organellar transcript abundance vastly exceeds nuclear transcription in plant cells (Havird and Sloan 2016; Forsythe et al. 2022). ERC signature between Clp and proteins likely involved in RNA processing may point to candidate proteins that could mediate crosstalk between plastid gene expression and plastid proteostasis status.

RNase E/G-like (RNE; AT2G04270) is a single-copy gene in *A. thaliana*. Its well-characterized homolog in *E. coli* is an endoribonuclease involved in processing pre-mRNAs and rRNAs. Likely because the targets of RNE are core gene expression components, it contributes to global gene expression patterns in *E. coli*. Furthermore, *E. coli* RNE is a core component and assembly factor of the ‘RNA degradosome’, a multiprotein complex involved in the degradation of bacterial mRNAs (Walter et al. 2010). In plants, RNE is a plastid-localized protein that shows similar enzymatic activity to the *E. coli in vitro* (Mudd et al. 2008; Schein et al. 2008). However, the specific *in vivo* role of RNE in plant plastids is less clear. RNE mutants display chloroplast phenotypes to different degrees (Mudd et al. 2008; Walter et al. 2010). Loss of RNE has been shown to lead to arrest of chloroplast development, loss of autotrophic growth, and reduced *psbA* and *rbcL* mRNA levels (Mudd et al. 2008), as well as reduced rRNA levels, leading to ribosome deficiency (Walter et al. 2010). It appears that RNE activity increases the stability of its targets by processing polycistronic RNAs to a mature state and potentially by removing polyadenylation signals that mark plastid mRNAs for degradation (Stoppel and Meurer 2012). Our results suggest that RNE is co-functional with plastid proteostasis machinery. Whether this apparent co-functionality stems from a specific unknown function of RNE or from the general relationship between plastid gene regulation and proteostasis remains an open question.

Our short-list contains two proteins with direct evidence of involvement in organellar RNA editing. First, White to Green 1 (WTG1; AT5G53080) is a tetratricopeptide repeat (TPR) protein, which contains tandem repeats of 34 amino acid motifs. TPR proteins (and related PPR proteins) play important roles in organelles (Lurin et al. 2004). While many PPRs are involved in RNA editing (Ma et al. 2017), TPR proteins, are typically associated with roles in organellar import (Mirus et al. 2009), gene expression (Hu et al. 2014), protein turnover (Park et al. 2007), and mediating protein-protein interactions (Blatch and Lässle 1999), all of which have been shown to be critical to organelle biogenesis (Hao et al. 2021). WTG1 is also required for chloroplast development (Ma et al. 2017). Unlike other TPR proteins, it appears to function in plastid RNA editing via interaction with two Multiple Organellar RNA Editing Factor (MORF) family proteins (Takenaka et al. 2012) in *A. thaliana* (Ma et al. 2017). WTG1 mutants were found to be deficient in editing in only two of the 43 plastid RNA editing sites (Ma et al. 2017). These results suggest that WTG1 could play a role in enhancing the function of the ‘RNA editosome’ a largely hypothetical complex whose composition is not fully resolved (Takenaka et al. 2013). If WTG1 plays such a role as an adaptor, this could position it to receive the brunt of the selective pressure exerted by hypothesized perturbations that occur in specific plant lineages during evolution (Forsythe et al. 2021). A conceivable example of such a rewiring event in RNA editing systems could be related to lineage-specific duplication and loss events of components of RNA editosome components, such as MORF proteins (Takenaka et al. 2013), in which case WTG1 may exhibit accelerated evolution in order to accommodate changes in the composition of binding partners.

A second member of our short-list with potential RNA editing activity is the uncharacterized PPR protein AT1G26460 (Lurin et al. 2004). Unlike the proteins described so far, AT1G26460 is localized in the mitochondrion. Little is known about the specific molecular function of AT1G26460, but it has been identified as one of three PPR proteins that exhibit significant co-expression with mitochondrial ribosomal protein S10 (RPS10; AT3G22300) (Doniwa et al. 2010). One of the other identified PPRs was further functionally characterized to be involved in mitochondrial RNA editing; however, the molecular function of AT1G26460 was not explored (Doniwa et al. 2010). Further, AT1G26460 was identified as one of 86 mitochondrial proteins with RNA binding activity (Bach-Pages et al. 2020). These findings provide tentative evidence that AT1G2646 may contribute to RNA editing in plant mitochondria.

The biggest mystery lies in why this N-mt protein exhibits ERC with plastid proteostasis systems. The answer may be tied to the intriguing finding that AT1G26460 protein expression was found to be upregulated in an *A. thaliana* mutant for ClpP2, the mitochondrial version of the Clp protease (Petereit et al. 2020). ClpP2 and ClpP1 are distant homologs, both originating from bacterial organelle progenitors; however, to our knowledge a functional or regulatory coordination has never been exhibited between the two systems. Our finding of an evolutionary connection of AT1G26460 to plastid ClpP1 combined with the regulatory connection between AT1G26460 and mitochondrial ClpP2 (Petereit 2020) raises the intriguing possibility that AT1G26460 could mediate a novel pathway of communication between plastid and mitochondrial proteostasis systems. The molecular mechanisms underlying coordination of plastid and mitochondria biogenesis are largely unknown; however, the ubiquitin proteosome system has been suggested as a potential coordination mechanism (He et al. 2023). Our results suggest a second potential mechanism involving PPR-mediated RNA editing and Clp proteases by which plastids and mitochondria coordinate their function.

### A possible retrograde plastid signaling mechanism mediated by nuclear-localized actin-related proteins

Our short-list of ERC hits contains two actin-related proteins, ARP5 (AT3G12380) and ARP7 (AT3G60830). These ERC hits represent examples of identified proteins that are not plastid localized. Instead, both ARP5 and ARP7 have been shown to localize to the nucleus in plants (Kandasamy et al. 2003, 2009). Both proteins have been implicated to be involved in epigenetic gene regulation, histone modification/recruitment, and DNA damage repair in yeast, mammals, and plants (McKinney et al. 2002; Kandasamy et al. 2003, 2005, 2009). In plants, ARP5 and ARP7 mutants are ubiquitously expressed in all cell types and knockout mutants are viable but display moderate to severe growth defects. In yeast and animals, the function of ARPs appears to be tied to interaction with the INO80 chromatin remodeling complex (Eustermann et al. 2018); however, in plants there is some evidence that ARP5 functions independently of the INO80 complex (Kandasamy et al. 2009).

Our finding that ARP5 and ARP7 exhibit strong signatures of ERC with plastid proteostasis machinery is somewhat surprising, given that their annotated functions take place in the nucleus. In general, actin filaments are known to physically contact plastids and are critical for the movement of chloroplasts within the cytoplasm in response to changing light conditions (Kadota et al. 2009). The physical interaction of actin with plastids appears to be mediated by proteins on the plastid protein import apparatus (Jouhet and Gray 2009), which is intriguing, given our finding that protein import is tightly linked to plastid proteostasis (see above). However, actin-related proteins are distantly diverged from conventional actin proteins, often displaying <50% sequence identity (McKinney et al. 2002). It is possible that ARP5 and ARP7 serve undiscovered moonlighting roles in nucleating actin filament assembly/branching in the cytoplasm to assist in light-dependent chloroplast movement. However, given the proven roles of ARP5 and ARP7 in chromatin remodeling, we speculate that instead ARP5 and ARP7 may represent long sought after components in plastid-to-nucleus (i.e. retrograde) signaling pathways.

A genetic screen yielded a series of genome-uncoupled (GUN) mutants that do not display characteristic coupling of nuclear-encoded and plastid-encoded photosynthesis genes (Susek et al. 1993). However, in the decades since that initial genetic screen, the molecular mechanisms underlying retrograde signaling remain unresolved (Tang et al. 2024). There is growing evidence that heme serves as a critical mobile retrograde signaling molecule (Woodson et al. 2011). Under this model, heme synthesized in the plastids, is released into the cytoplasm and binds heme-sensitive enzymes, including enzymes involved in regulating nuclear gene expression, allowing for the photosynthesis status of plastids to be communicated to the nucleus. Heme-dependent signaling in response to photosynthetic perturbation is the most widely recognized form of retrograde signaling; however, it has been suggested that retrograde signaling in response plastid-proteostasis as part of the plastid unfolded protein response (UPR) is another underappreciated and important form of retrograde signaling (Richter et al. 2023). For example, when plastid Clp genes are mutated or inhibited, nuclear-encoded proteins involved in protein folding/degradation are upregulated (Sjögren et al. 2004; Zybailov et al. 2009; Perlaza et al. 2019), presumably via retrograde signaling. In green algae, this upregulation appears to be mediated by a cytoplasmic kinase, MARS1, representing the first identified UPR retrograde signaling factor (Perlaza et al. 2019). However, the full set of proteins involved in this newly identified pathway, including those that physically localize to nuclei to directly impact nuclear gene expression, are not known. Furthermore, while there is evidence of UPR retrograde signaling spanning all plants, a functional ortholog of the green algal MARS1 has not been identified in higher plants (Perlaza et al. 2019). Based on our results, we hypothesize that ARP5 and/or ARP7 could play a previously undiscovered role in UPR retrograde signaling by affecting chromatin remodeling to elicit a UPR nuclear expression program in response to perturbation of plastid proteostasis. We expect that this hypothesis (and the others presented above) will motivate experiments to understand cross-compartment connectivity in plant proteostasis systems. Such work will have to potential to reveal important cellular mechanisms, while also adding important datapoints to the discussion of the types of functional interactions that underly signatures of ERC.

## Methods

### Obtaining genomic datasets

Our study analyzed the dataset that was previously used to probe plastid-nuclear interactions in specific angiosperm taxa that were sampled to include known plastome acceleration events (Forsythe et al. 2021). Sampling was designed to capture diverse angiosperm lineages and avoid close relatives. This approach was taken to achieve a ‘star phylogeny’ in which most of the evolution takes place along the external branches of the tree, allowing us to employ the ‘root-to-tip’ method of measuring branch length by minimizing bias associated with phylogenetic non-independence (Gatts et al. 2024). We analyzed our sampled proteomes in *Orthofinder* (Emms and Kelly 2015, 2019) and used the *Orthofinder* results as input for out downstream ERC analyses. We included two outgroups (*Amborella trichopoda* and *Liriodendron chinense*) and treated the remaining species as our ingroup during ERC analyses.

### All-by-all Evolutionary Rate Covariation analyses

This study builds on prior one-by-all ERC analyses to perform all-by-all analyses using the newly developed *ERCnet* v1.1.0 program (Gatts et al. 2024). Analyses were run on a linux server, using 48 threads for parallel computing. The full *ERCnet* run took 10.7 hours. The four steps of the *ERCnet* workflow were run with the following commands. See the *ERCnet* GitHub page (https://github.com/EvanForsythe/ERCnet) for detailed descriptions of each parameter.

`./Phylogenomics.py -j <name of job> -o <path to orthofinder results> -x <path to RaxML installation> -m 48 -s -p 3 -r 4 -T <path to Julia installation> -n 2`

`./GTST_reconciliation.py -j <name of job>`

`./ERC_analyses.py -j <name of job> -m 48 -s Atha -b R2T`

`./Network_analyses.py -j <name of job> -m R2T -c both -p 0.0001 -r 0.5 -y fg`

`-Func_cat -s Atha -f <name of ERC analyses results file>`

### Network visualization and enrichment analyses

Networks were visualized and analyzed with *Cytoscape* (Shannon et al. 2003), making use of the *GraphML*-formatted file output by *ERCnet*. Edges were weighted by the strength of correlation, *Cytoscape’s* Select>Nodes>First neighbors of selected nodes option was used to obtain subnetworks surrounding focal nodes of interest. We used *Cytoscape’s* ClusterMaker cluster network> Community cluster (GLay) option to find the cluster containing plastid partition nodes. We used *Cytoscape BINGO* (Maere et al. 2005) plugin to perform GO functional enrichment analyses (Ashburner et al. 2000).

### Statistical tests of functional clustering

The *--Func_cat* feature of *ERCnet* (implemented in *Network_analyses*.*py*) was used to statistically assess clustering of functionally related proteins across the network using the assortativity coefficient summary statistic from *igraph* (Csardi and Nepusz 2006). We applied these tests using two different types of functional categories, (1) subcellular localization of proteins as predicted by *CyMIRA* (Forsythe et al. 2019) and a genome-wide catalog of proteostasis genes (Majsec et al. 2017). For both assortativity analyses, we generated a null distribution by randomizing the attribute assignments across the network and calculating the assortativity coefficient. We replicated the random-attribute networks 1000 times to create a null distribution, which were centered around assortativity coefficient = 0. We used null distributions to perform two-tailed z-tests to ask whether the observed assortativity coefficients significantly differ from zero.

## Data Availability

*ERCnet* output used in this analysis have been deposited at Dryad Digital Repository and can be accessed at: https://doi.org/10.5061/dryad.hx3ffbgp7

## Acknowledgements

This work was supported by a National Science Foundation (NSF) grant (IOS-2114641).

